# Predicting neural responses to intra- and extra-cranial electric brain stimulation by means of the reciprocity theorem

**DOI:** 10.1101/2024.08.04.603691

**Authors:** Torbjørn V. Ness, Christof Koch, Gaute T. Einevoll

## Abstract

Electrical stimulation of nervous tissue is grounded in well-established biophysical principles, yet understanding its precise effect on neural activity remains challenging. This represents a major obstacle to both scientific and clinical applications of electrical brain stimulation.

We here show that the reciprocity theorem of electromagnetism can be applied more broadly than previously acknowledged, providing a framework for elucidating the effects of electrical brain stimulation on neurons. Through this new perspective, we account for the observed weak frequency-dependence of extracellular electrical stimulation–induced membrane potential changes, and demonstrate a 1/*r* falloff for nearby and 1/*r*^2^ falloff for distant neurons.

We also show that for transcranial electrical stimulation, the susceptibility of a neuron to the stimulation is directly proportional to the size of the current dipole moment resulting from somatic current input. The induced somatic membrane potential changes are small and exhibit weak spatial but strong orientation dependency across human neocortex.

The reciprocity-based approach to electrical stimulation reproduces a large body of previous experimental data, and provides new insights into how ES affects neurons.

## 1. Introduction

The reciprocity theorem (RT) is a foundational principle in physics, stating that in linear systems the source and the measurement sites can be swapped without a change in the measured signal. It was originally formulated more than 150 years ago in the context of electromagnetism [Helmholtz, 1853], but has extended to a wide range of disciplines [Malmivuo, 2010], including antenna design [Hoop and Jong, 1974], biomedical imaging [Malmivuo, 1976], earthquake seismology [Wapenaar et al., 2019] and distance measurements in cosmology [Renzi et al., 2022]. In the context of neuroscience, the RT has been used to swap the location of extracellular current sources— sometimes in the form of equivalent current dipoles—and extracellularly measured electric potentials [Plonsey, 1963; Rush and Driscoll, 1969]. Here we draw attention to the fact that the RT is also valid for intracellular current sources in the approximately linear (subthreshold) domain. This opens up for entirely new applications of the RT, namely to directly study how the somatic membrane potential is affected by extracellular current stimulation in a straightforward manner.

Extracellular potentials in neural tissue can be generated by exogenous currents, delivered either directly into extracellular tissue via invasive electrodes, referred to here as extracellular electric stimulation (ES) [Sironi et al., 2011; Histed et al., 2013], or in a non-invasive manner, via transcranial electric stimulation outside the head (tES), which includes tACS, tDCS, and temporal interference (TI) [Liu et al., 2018]. Medicine and neuroscience have more than two centuries of experience with ES and tES [Sironi et al., 2011; Vincent et al., 2016], mapping the function of brain areas [Vincent et al., 2016], studying functional connectivity [Matsumoto et al., 2007; Rossel et al., 2023], guiding awake neurosurgery [Duffau, 2005] and treating neurological disorders such as Parkinson’s disease, dystonia, depression, and epilepsy [Sironi et al., 2011; Cagnan et al., 2019; Vissani et al., 2020].

We have a reasonably good understanding of the biophysics underlying single neurons and how these generate extracellular potentials *V*_e_, stemming from membrane currents propagating through an essentially ohmic extracellular medium [Nunez and Srinivasan, 2006; Ilmoniemi and Sarvas, 2019; Halnes et al., 2024]. Computer modeling of these mechanisms, based on the low-frequency, electro-quasistatic, ohmic-current-dominated approximation to the Maxwell equations, to study neural function is a mature field that has received substantial attention over the past decades [Rall and Shepherd, 1968; Holt and Koch, 1999; Nunez and Srinivasan, 2006; Buzsáki et al., 2012; Einevoll et al., 2013; Ilmoniemi and Sarvas, 2019; Ness et al., 2022; Halnes et al., 2024].

The opposite situation, that is, how extracellular potentials influence sub-threshold and spiking activity is, on the other hand, insufficiently understood. This is true for intracranial microstimulation [Histed et al., 2009; Aberra et al., 2018; Rossel et al., 2023; Lee et al., 2024], but even more so for ES that targets larger brain regions, like Deep Brain Stimulation (DBS) [Sironi et al., 2011; Ashkan et al., 2017; Vissani et al., 2020]. For tES, and, in particular, the emerging technique of TI [Mirzakhalili et al., 2020; Violante et al., 2023; Vieira et al., 2024; Caldas-Martinez et al., 2024], the causal mechanisms underlying any behavioral effects, above and beyond indirect effects (e.g., peripheral stimulation), remain controversial [Liu et al., 2018; Beliaeva et al., 2021].

Given the known biophysics underlying ES in the low kHz range [McNeal, 1976; Rattay, 1986; McIntyre and Grill, 2000; Joucla and Yvert, 2012; Tveito et al., 2017; Krause et al., 2023], it is, in principle, straightforward to estimate the effect of ES via computational modeling [McIntyre et al., 2002; Miocinovic et al., 2006; Aspart et al., 2018; Aberra et al., 2018; Mirzakhalili et al., 2020; Aberra et al., 2023]. However, given the daunting complexity of nervous tissue, together with the multi-dimensional nature of ES (geometry of stimulating electrodes relative to the distance and morphology of neurons, temporal-frequency content, amplitude etc.), translating individual simulations into an intuitive understanding of how ES affects neurons has proven challenging.

This difficulty is well illustrated by the recent results of Lee et al. [2024], where even single-cell subthreshold—and therefore presumably approximately linear—effects of ES in cortical slices are surprising and hard to understand: While neurons have cell-type specific and highly variable frequency-dependent electrical properties, Lee et al. [2024] observed robust frequency-independent ES responses that were similar across cell-type, brain region, and even species.

This is where the *Reciprocity Theorem* (RT) can come to the rescue. It applies to Maxwell’s equations of electromagnetism for time-invariant, linear media, in which the current is linearly related to the electric field. Intuitively, reciprocity implies that the relationship between an oscillating current and the resulting electric field is unchanged if the points where the current is placed and where the field is measured are interchanged [Helmholtz, 1853; Plonsey, 1963; Rush and Driscoll, 1969]. The RT has often been applied when calculating extracellular potentials from current sources [Moffitt and McIntyre, 2005; Lempka et al., 2011; Ness et al., 2015; Tharayil et al., 2024], or EEG signals from current dipoles [Ziegler et al., 2014; Huang et al., 2016; Dmochowski et al., 2017; Halnes et al., 2024; Tharayil et al., 2024].

The RT is, however, also applicable if the current source is intracellular, with far-reaching consequences. In our context, the RT states that for any tissue characterized by an electrical network consisting of ideal capacitances, inductances, resistances, and batteries, a change in the membrane potential *V*_m_(**r**′) due to an extracellular current source *I*_stim_ at location **r** is equivalent to the change in the extracellular potential *V*_e_(**r**) due to a current input *I*_stim_ at cellular location **r**′ (Figure 1). This is convenient because it allows us to apply our understanding of how extracellular potentials are generated by neural activity [Halnes et al., 2024] to better understand the effect of ES on the membrane potential of single neurons.

**Figure 1:**
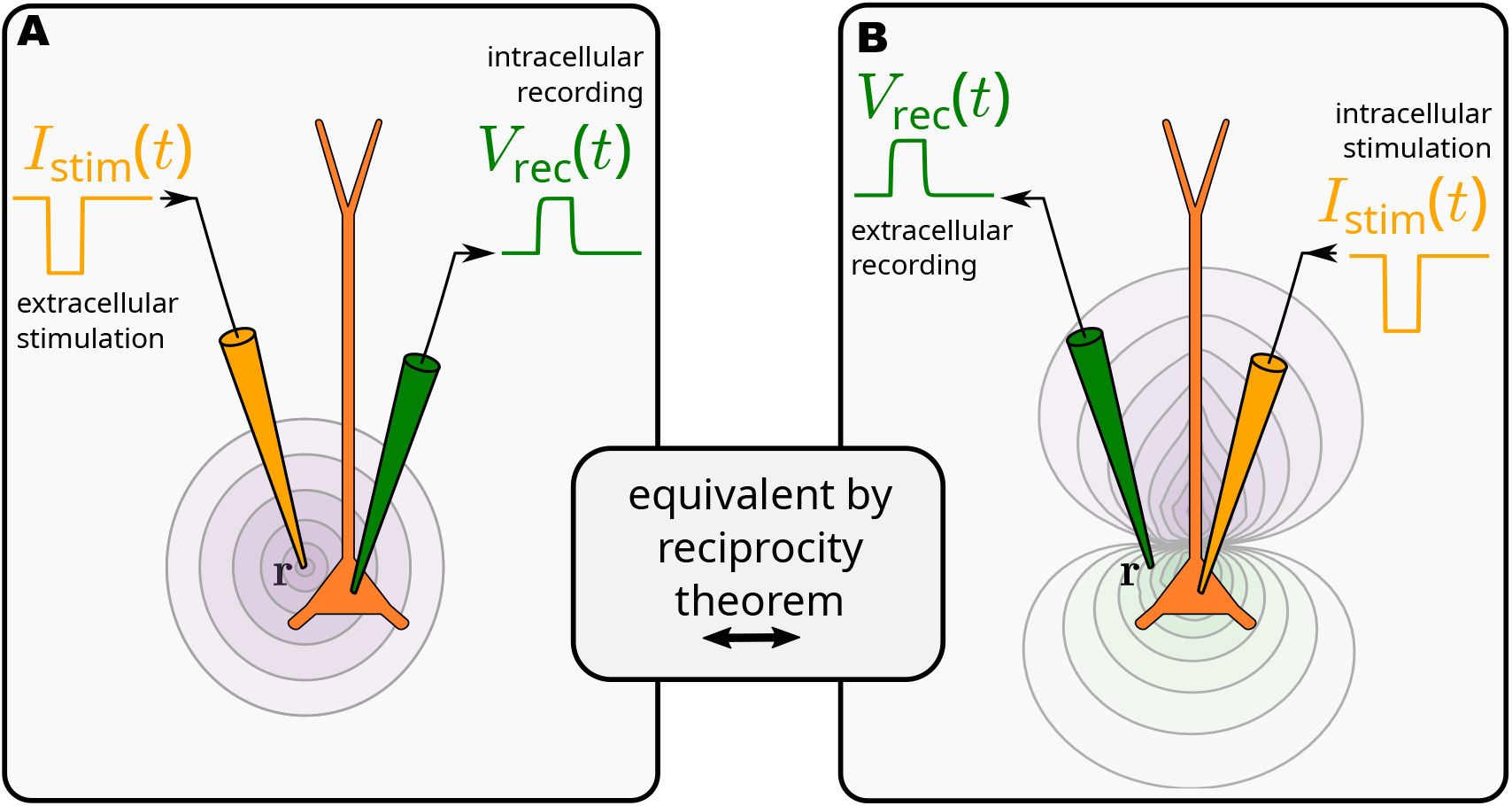
The reciprocity theorem in the context of electrical brain stimulation. The strategy is to use the situation on the right, in which an injected intracellular current *I*_stim_ induces a change in extracellular potential, *V*_rec_ at location **r**, to infer what happens on the left, in which the same current is injected at **r**, inducing the identical membrane potential *V*_rec_. Equally, the setup on the left can be used to infer the situation on the right.

The RT is useful in systems with locations of special interest where one would like to know the measured signal resulting from sources at arbitrary locations. The cell body of a neuron is such a singular place. In the traditional approach to simulating ES, we simultaneously calculate the *V*_m_ response at all cellular locations in response to an extracellular current source. However, experimentalists are often primarily interested in the somatic response to ES, both because this is by far the most accessible region of a neuron, as well because the soma is electrically tightly coupled to the axon hillock where *V*_m_ is transduced into action potentials that are the relevant output of any spiking neuron. That is, knowing the somatic *V*_m_ is highly informative about the likelihood of that neuron generating one or more action potentials.

Let us compare the two approaches.

The standard framework has two steps; the first one involves calculating the extracellular potential resulting from the ES. The second step is simulating the effect on *V*_m_ from the *spatial gradient* of this extracellular potential over the neural morphology. Both these steps can be numerically evaluated, but building a good, intuitive understanding is hard. In particular, the second step is opaque, and non-intuitive, even to long-time practitioners of the art, so that insights do not transfer well to other scenarios or parameters.

With intracellular somatic current input, the situation is easier - calculating the resulting membrane currents, and the resultant *V*_e_ everywhere has been the goal of cable theory for the past century thanks to the efforts of Hodgkin, Huxley, Cole, Rall, Jack, Noble, Tsien and others, with a well-developed calculus and intuition [Jack et al., 1975; Koch, 1999; Halnes et al., 2024]. From a somatic intracellular current source, we simultaneously compute *V*_e_ at all relevant extracellular locations. Through the RT, we therefore know the somatic *V*_m_ response to ES from any of these locations. The RT-based approach therefore allows us to take a *soma-centric viewpoint* that is advantageous.

The RT assumes linearity, and since neurons often behave approximately linearly in the sub-threshold regime [Bikson et al., 2004; Tran et al., 2022; Gaugain et al., 2024] (as we will demonstrate later), the RT can be applied to study the subthreshold effects of ES. Thus, if we can calculate the extracellular potential resulting from somatic current input, we know the somatic membrane potential response to the same extracellular current (Figure 1). Perhaps counterintuitively, the RT is also, as demonstrated here, germane to tES as practiced today, where ES current flow is limited to 4 mA or less [Antal et al., 2017].

We here first show how the RT is applicable to accurate simulation of ES on biophysical-detailed and simplified models of cortical neurons. We demonstrate how this approach leads to a better understanding of ES on different neural elements, and how this can explain many empirical findings, and make new predictions. We then apply the RT to infer the effect of tES on single neurons, deep inside the human brain.

## 2. Results

### 2.1. Validating the reciprocity-based approach

We consider the effect of an extracellular sinusoidal current *I*_stim_ at location **r**_EC_ on the somatic *V*_m_ of a canonical rat cortical layer 5 pyramidal cell with detailed dendritic morphology and ten voltage- and calcium-dependent membrane conductances distributed through the cell (Figure 2A) [Hay et al., 2011].

**Figure 2:**
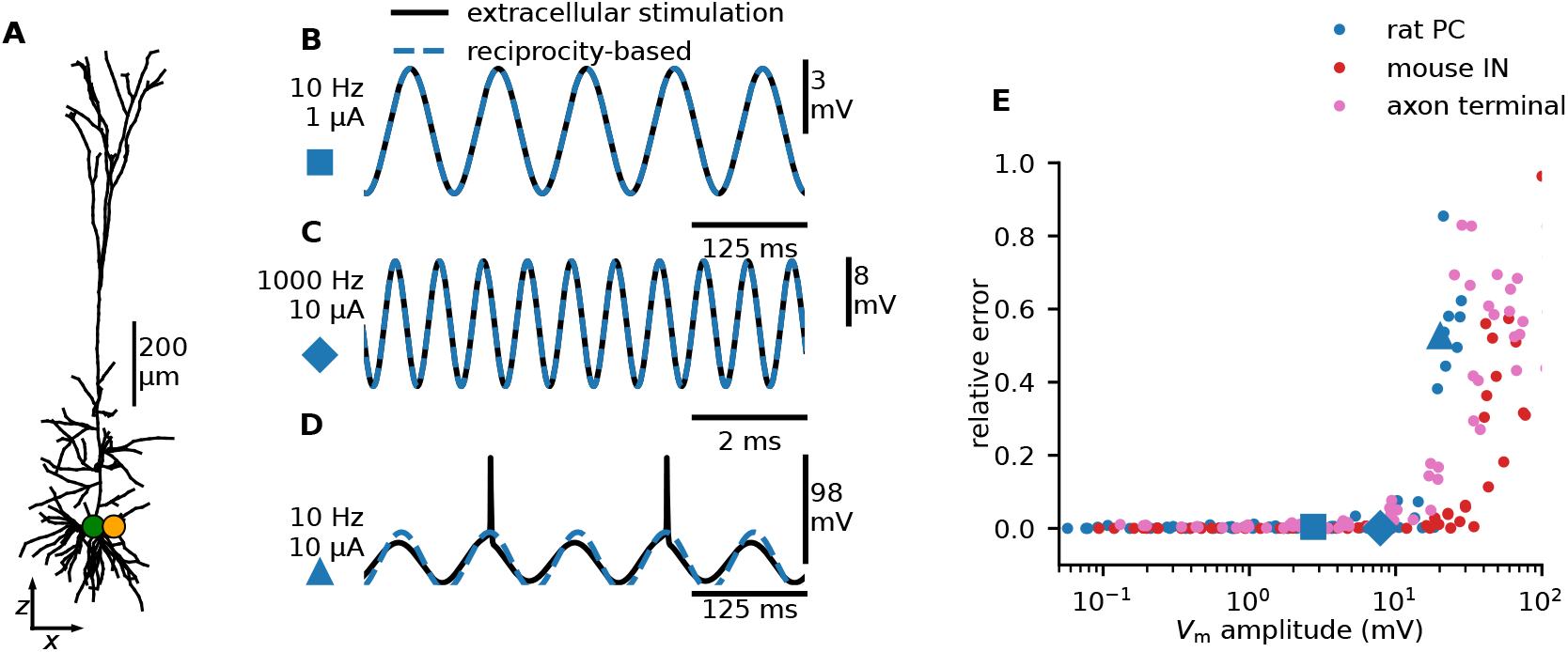
Applying the reciprocity theorem to subthreshold ES of a compartmental model of a cortical neuron. **A:** Stimulating a canonical cortical layer 5 pyramidal cell model [Hay et al., 2011] with a nearby current (orange dot, 50 µm outside the soma), while recording the somatic membrane potential (green dot). **B:** *V*_m_ responding to ES of 10 Hz and 1 µA (black line). The reciprocal situation, *i.e*., injecting the same current into the soma while measuring changes in *V*_*e*_ at the orange dot, leads to indistinguishable results (blue dashed line). Note that the reciprocity-based solution predicts the membrane potential response, that is, the deviation from the baseline, but not the resting membrane potential itself. **C:** Same as in B, but for a 1000 Hz, 10 µA current. **D:** Same as in B, but for a 10 Hz, 10 µA current. This combination of amplitude and stimulation frequency evokes action potentials, and is therefore outside the linear regime. **E:** The relative error as a function of *V*_m_ at the stimulation frequency, for three different models: the layer 5 pyramidal cell from panel A, a mouse cortical layer 5 GABAergic PV interneuron, and a straight, unbranched myelinated axon with periodic nodes of Ranvier (see Methods). We injected different currents (0.1, 0.5, 1, 5, 10, 50, 100 µA) at different frequencies (1, 10, 100, 1000 Hz) and distances from the soma (25, 50, 100 µm). The instances of panels B, C, and D, are marked, respectively, by a large blue square, diamond, and triangle. The relative error is defined as SD(*V*_ES_ − *V*_reciprocity-based_)/SD(*V*_ES_).

In accordance with the RT, the model shows that *V*_m_ induced by ES—here a current of 1 µA and 10 Hz induced a peak *V*_m_ of 2.7 mV (Figure 2B)—is indistinguishable from *V*_e_ at location **r**_EC_ due to intracellular current input *I*_stim_ to the soma (Figure 2B,C, “reciprocity-based”).

Amplitudes for ES are, however, typically large compared to intracellular currents (mA vs. nA) which would trigger hyper-excitability if delivered intracellularly to active cell models. To readily apply the RT to active cell models, we avoided this methodological issue in Figure 2 by using a fixed subthreshold current amplitude for the intracellular current to the active cell model, and rescaling *V*_e_ to reflect the current amplitude used for the ES. That is, while ES is applied directly to the fully active cell model, the reciprocity-based approach is a linear extrapolation of the subthreshold response of the active cell model (see Methods).

Indeed, the active cell model effectively responds linearly to ES, in line with experimental observations [Bikson et al., 2004; Gaugain et al., 2024]. This remains true if ES is increased to 10 µA with a frequency of 1 kHz (Figure 2C), giving a peak *V*_m_ of 7.8 mV from rest (Figure 2C).

However, the closer *V*_m_ veers to the cell threshold, the less applicable the RT will be. Thus, for ES that directly evokes spikes (here a current of 10 µA at 10 Hz), the RT approach is not reliable (Figure 2D). We quantify the relative error of applying the reciprocity-based approach. This error is zero for passive cell models, a few percent for active models if the evoked *V*_m_ *<* a few mV (Figure 2E, dots) but abruptly increase for ES amplitudes that are driving the values of *V*_m_ close to or above the threshold for spiking.

### 2.2. RT for studying the effects of ES on different neural elements

The effect of ES on the somatic *V*_m_ is informative about how the neuron’s spiking activity will be affected (while spike initiation happens at the axon initial segment, it is electrically closely coupled to the soma). Via the RT this can be studied by considering *V*_e_ from a somatic current input. This is convenient because we have a good understanding of how neurons generate extracellular potentials [Holt and Koch, 1999; Halnes et al., 2024]. We therefore review some general features of extracellular potentials generated by current input to neurons, and consider the implications for how neurons are affected by ES.

For clarity of exposition, we restrict ourselves to passive cell models, but as demonstrated in Figure 2, our results are directly relevant for active cell models operating in the subthreshold regime.

#### 2.2.1. Pyramidal cells

We consider a passive version of the cortical rat layer V pyramidal cell model from Hay et al. [2011] and calculate *V*_e_ resulting from somatic white-noise current input. All frequency components of the white-noise current have an equal amplitude of 1 nA (Figure 3A).

**Figure 3:**
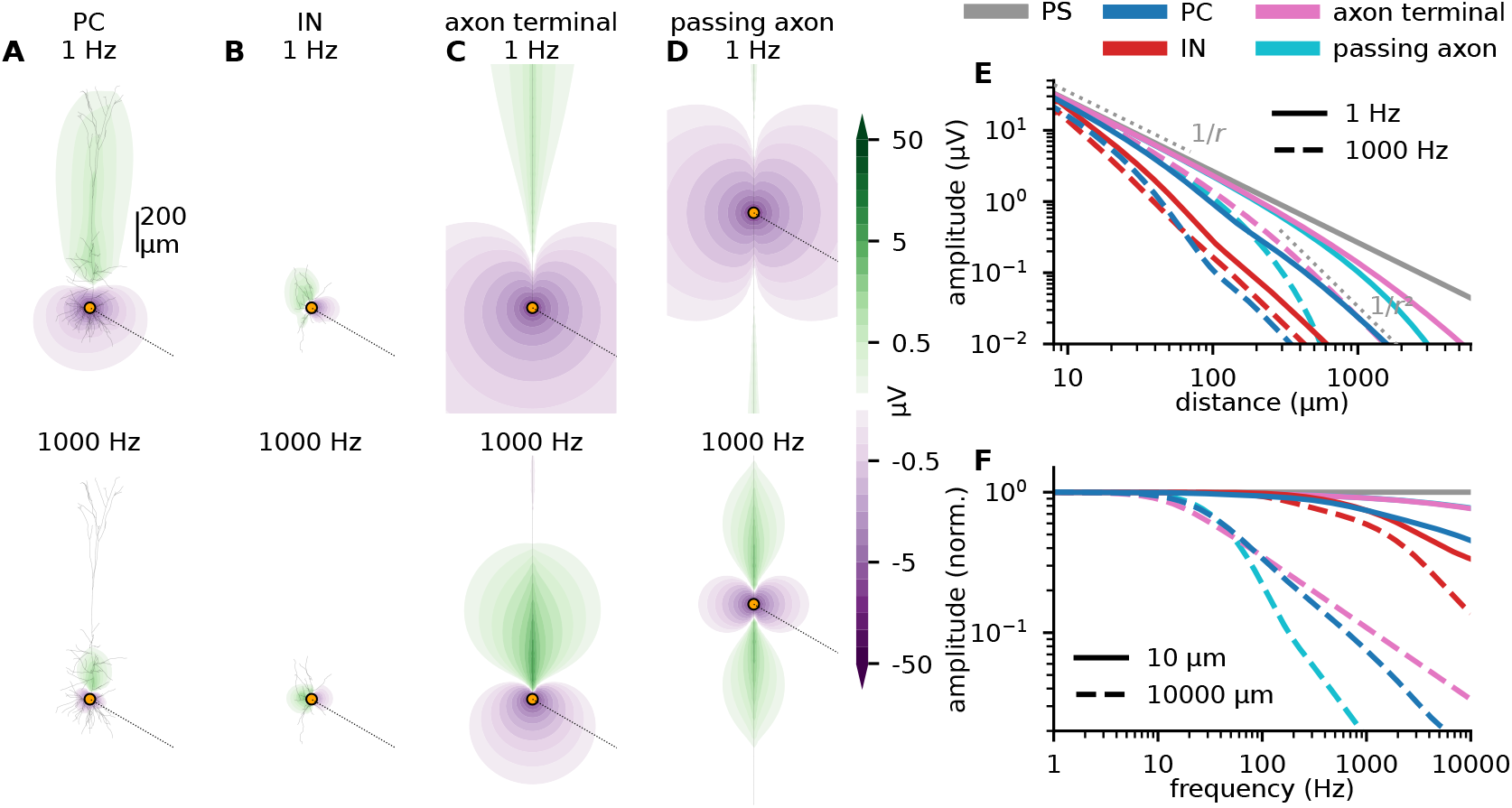
Extracellular potentials surrounding different neural models are informative about the effect of ES. **A:** *V*_e_ following somatic white-noise current input, where each frequency component has an amplitude of 1 nA (see Methods), to a passive rat cortical layer 5 pyramidal cell model (passive version of the model shown in Figure 2A). *V*_e_ is calculated on a dense grid around the neuron. Through Fourier analysis, the 1 Hz (top) and 1000 Hz (bottom) components of *V*_e_ are extracted (see Methods). **B:** Same as in panel A, except for a passive mouse cortical layer 5 interneuron model. **C:** Same as in panel B, except for a passive myelinated axon model, with the current input at the terminal end. The axon is oriented vertically, parallel to the apical dendrite in panel A. **D:** Same as in C, except that the current input is in the middle of the axon. **E:** *V*_e_ as a function of distance from the site of current input, calculated along the dotted lines in A-D (at -30^°^ relative to the horizontal direction). Different colors correspond to the four different scenarios in panels A-D, as well as *V*_e_ from a point source (PS). Full and dashed lines correspond to 1 Hz and 1000 Hz input respectively. **F:** *V*_e_ as a function of frequency for white noise current input, calculated at two different distances along the dotted lines in A-D. Full and dashed lines correspond to 10 µm and 10,000 µm distance respectively. All amplitudes are normalized to the value at 1 Hz.

The amplitude of a frequency component of *V*_e_ at any given location corresponds, through the RT, to the amplitude of *V*_m_ caused by a sinusoidal ES at that particular frequency and location. Lee et al. [2024] recorded somatic *V*_m_ responses to ES of 100 nA, at a distance of 50 µm away from the soma (in the frequency range 1–100 Hz) of about 0.3 mV. How does this compare to our results?

At 1 Hz and a distance of 50 µm, a 1 nA current amplitude yields a *V*_e_ of about 3 µV (Figure 3E). Extrapolating linearly, the predicted *V*_m_ of an ES of 100 nA will be 0.3 mV, very similar to Lee et al. [2024]. This serves as a qualitative check, confirming that our results are biologically plausible. Note that we have effectively quantified the subthreshold ES response of the neuron model for any ES location through a single simulation with *V*_e_ calculation. For the traditional approach to simulating ES, each ES location requires a separate simulation.

The *V*_e_ from somatic current input has a dipole-like shape, aligned with the orientation of the apical dendrite, perpendicular to the cortical surface (Figure 3A), as expected from theory [Halnes et al., 2024]. The negative regions of *V*_e_ correspond to where *V*_m_ is 180 degrees “out of phase” with the ES, that is, where *V*_m_ will be positive (depolarizing) in response to the negative phase of ES. This is again consistent with literature [Lafon et al., 2017; Lee et al., 2024]. ES located in the crossover region between the negative and the positive regions of *V*_e_ will have a negligible effect on the somatic *V*_m_, while for ES in the positive region outside the apical dendrite, the somatic *V*_m_ will “be in phase” with the ES, that is, the *V*_m_ response will be positive to the positive phase of ES.

ES will always depolarize one region of a neuron, while hyperpolarizing others [Basser and Roth, 2000; Radman et al., 2009; Lafon et al., 2017; Aberra et al., 2023]. Indeed, it is impossible to devise an ES that will have a purely depolarizing or hyperpolarizing effect across an entire neuron. This can also be shown through the RT: Current conservation ensures that at any given time, a neuron’s membrane currents must sum to zero, and extracellular potentials are therefore intrinsically dipole-like in nature, with both positive and negative regions. A current input to the soma or to the apical dendrite will give rise to dipole-like extracellular potentials of opposite orientation [Lindén et al., 2010; Næss et al., 2021; Halnes et al., 2024]. As a consequence, ES will have opposite effects on somatic and apical dendritic *V*_m_.

Interestingly, in analogy to how pyramidal neurons are expected to dominate LFP/EEG/MEG signals because they are spatially aligned [Lindén et al., 2011; Buzsáki et al., 2012; Halnes et al., 2024], this geometrical alignment also predicts that pyramidal cells with approximately the same orientation and distance from an extracellular current source will be affected in the same manner. We can therefore expect distant ES to have a synchronizing effect on pyramidal neurons next to each other with the same orientation (*i.e*., in the same gyrus; [Aberra et al., 2023]).

The *V*_e_ from somatic current input becomes more spatially confined at higher frequencies (Figure 3A), as expected from intrinsic dendritic filtering [Lindén et al., 2010; Halnes et al., 2024]. This suggests that, in general, low-frequency ES will have a stronger effect on *V*_m_ than high-frequency ES. Importantly, however, *V*_e_ close to the soma is relatively frequency-independent below about 100 Hz (Figure 3F). This surprising result—the effect of ES on near-by cell bodies is approximately frequency independent—was reported—but left unexplained—in experimental ES work by Lee et al. [2024].

As earlier noted, in the traditional approach to simulating ES, each ES location requires a separate simulation, which has made general insights regarding the distance-dependence of ES hard to extract. Through the RT we can, however, draw from the well-established theory of extracellular potentials, and how they, for example, decay with distance from the electrode. To understand how ES decays with distance, assuming an infinite, homogeneous and isotropic volume conductor, we approximate *V*_e_ at location **r** for a model with *N* compartments via the point-source approximation [Holt and Koch, 1999; Halnes et al., 2024],

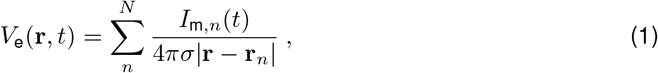

where *I*_m,*n*_ is the membrane currents of compartment *n, σ* is the extracellular conductivity, and **r**_*n*_ is the position of compartment *n*.

Sufficiently close to a soma (at **r**_*s*_=0), *V*_e_ will be dominated by the somatic membrane current alone [Pettersen and Einevoll, 2008; Halnes et al., 2024],

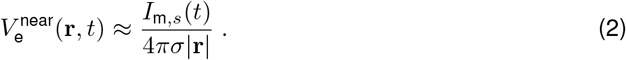

This “monopole”-like approximation is accurate as long as |**r**| *<<* the frequency-dependent length constant [Halnes et al., 2024], and the region where *V*_e_ ∝ 1*/*|**r**| is valid can be roughly estimated from Figure 3E. In absence of other inputs, the somatic membrane current itself will be dominated by the input current which counts as part of the membrane current *I*_m,*s*_ ≈ *I*_stim_,

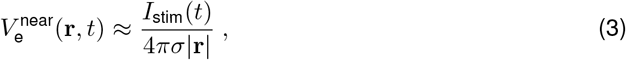

confirmed in Figure 3E (blue versus gray line). It follows that in the near-field limit, the amplitude of *V*_e_ will be approximately frequency-independent, and that for near-by ES, the specific membrane properties are of limited importance for determining the response of *V*_m_. The dominating factor is the ES current itself, as seen in our simulations for different cell models (Figure 3E).

Sufficiently far away from the neuron, we can express *V*_e_ in terms of the current-dipole approximation [Halnes et al., 2024],

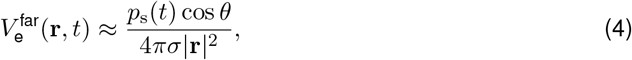

where *p*_s_ is the current-dipole moment of the cell resulting from the somatic current injection, and *θ* is the angle between the dipole orientation and the measurement point. In this far-field regime, *V*_e_ decays with distance as 1*/r*^2^ (“dipole”-like) [Pettersen and Einevoll, 2008; Halnes et al., 2024], and has a clear, albeit still relatively weak, frequency dependence above about 10 Hz (Figure 3F). From use of the RT, these properties are expected to apply to responses of *V*_m_ to distant ES as well.

Interestingly, we deduce that for distant ES—like tES—the somatic *V*_m_ response of a neuron is proportional to the current dipole moment of that neuron in response to somatic current input *p*_s_. Cellular features which maximize *p*_s_ will therefore also maximize the ES sensitivity. We will return to this point.

#### 2.2.2. Interneurons

Somatic current to a mouse layer V PV-interneuron model results in a *V*_e_ with a dipole-like shape (Figure 3B), whose orientation depends on the geometry of dendrites. Since interneurons lack apical dendrites, the effect of ES on a population of interneurons will be more variable than for pyramidal neurons, and we cannot *a priori* expect the ES to have a synchronizing effect [Lindén et al., 2010; Næss et al., 2021; Aberra et al., 2023].

Lee et al. [2024] reported *V*_m_ of similar amplitudes in pyramidal neurons and interneurons responding to nearby ES. This is consistent with our observations (Figure 3E). However, at least for low frequencies, *V*_e_ from interneurons enter the far-field regime at distances closer to the soma than for pyramidal neurons [Pettersen and Einevoll, 2008]. Thus, in this regime, ES on pyramidal cells will be stronger than on inhibitory neurons for ES that is more than some tens of micrometers away from the somas (Figure 3E).

*V*_e_ for the interneuron is almost independent of frequency below 1000 Hz (Figure 3F), as it is too compact for intrinsic dendritic filtering to have any low-pass filtering effect [Lindén et al., 2010]. These results indicate that for distant (e.g. transcranial) ES, low-frequency stimulation will have a predominately excitatory effect (Figure 3E, solid lines, blue versus red), while high-frequency ES will be more balanced, or even biased towards having an inhibitory effect (Figure 3E, dashed lines, blue versus red), in line with experiments [Caldas-Martinez et al., 2024].

#### 2.2.3. Axons

Next, we consider a current input to an axon end or terminal (Figure 2E). While this is difficult to achieve in practice due to their minute size, this is irrelevant for the validity of the RT.

*V*_e_ resulting from the current input has a dipole-like shape (Figure 3C), falling of as 1*/r* close by, and as 1*/r*^2^ far away (Figure 2E). The amplitude of *V*_e_ is higher than for the same input to cell bodies (Figure 3E). According to the RT, the same should hold for the amplitude of the *V*_m_ response to ES, compatible with both experiment [Nowak and Bullier, 1998] and modeling [Aberra et al., 2018].

Interestingly, *V*_e_ following current input to the middle of the axon model (Figure 3D) gives rise to “quadrupole-like”, shape, with the effect of ES decaying as 1*/r*^3^ in the far-field [Halnes et al., 2024]. That is, axon terminals are more excitable to ES than passing axons, although the difference is only visible beyond a certain distance (Figure 3E).

### 2.3. Analytical investigation of ES in simple cell models

To understand complex physical systems, it helps to start with tractable models capturing the essential underlying principles. Such simple models for the effect of ES on neural activity have been lacking. Through the RT we can, however, draw from analytic solutions of *V*_e_ from somatic current input to a passive ball and stick model, and directly apply the resulting formulas to investigate how the neuron model is affected by subthreshold ES. Near the soma, *V*_e_ is dominated by the somatic membrane current alone (equation (2)), while sufficiently far away the current-dipole approximation holds (equation (4)). To derive an analytical expression for how a neuron is affected by ES we need expressions linking an arbitrary somatic input current to the resulting somatic *V*_m_ for the near-field case, and to the resulting current-dipole moment for the far-field case.

Pettersen et al. [2014] derived expressions for somatic membrane currents 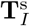, and the dipole moment 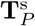. For an arbitrary input current **I**_in_ (for example, white noise), 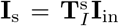 and 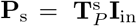 (boldface symbols signify complex numbers). Using these expressions (see Methods) confirms that *V*_e_ decays as 1*/r* in the near-field and as 1*/r*^2^ in the far-field regime (Figure 4A).

**Figure 4:**
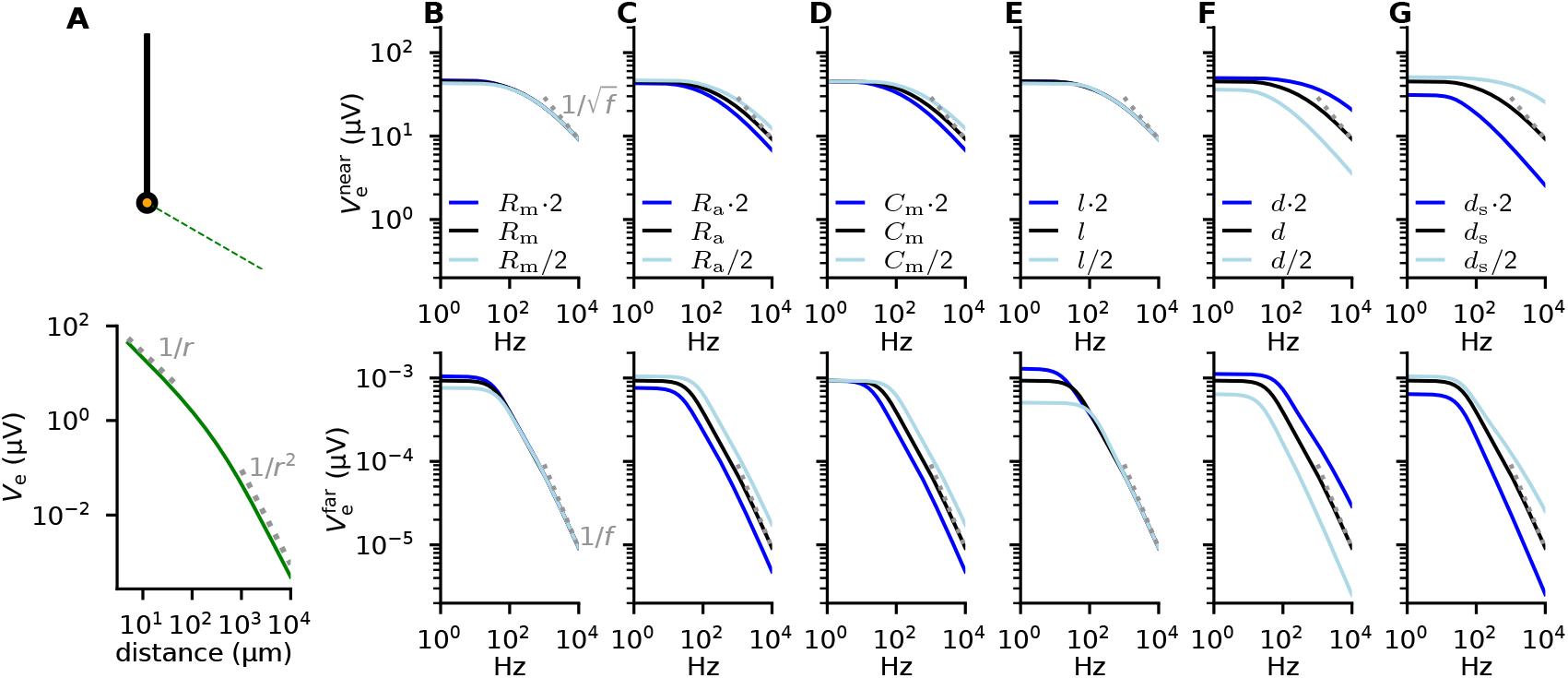
Analytical expressions for *V*_e_ gives insight into the parameter-dependence of ES. **A:** Illustration of how *V*_e_ decays with distance from the soma of a ball and stick neuron (see schematic) receiving a somatic current input with a frequency of 1 Hz and an amplitude of 1 nA. The bottom panel shows results from a numerical simulation of changes in *V*_e_ with increasing distance. Grey dotted lines indicate expected trends in the near (1/*r*) and far-field (1/*r*^2^) regimes. **B-G:** The amplitude of *V*_e_ as a function of frequency for a white-noise input current where each frequency component has an amplitude of 1 nA, and *V*_e_ is measured either very near (top) or very far away from (bottom) the soma. The three lines correspond to different parameter choices for a single parameter (columns), either default (black), increased by a factor of two (blue), or decreased by a factor of two (light blue). The parameters are membrane resistance *R*_m_ in B, axial resistance *R*_a_ in C, membrane capacitance *C*_m_ in D, length of the stick *l* in E, dendritic diameter *d* in F, and somatic diameter *d*_s_ in G. The grey dotted lines show the expected trends in the high-frequency limit in the near- 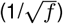 and far-field (1/*f*) [Pettersen et al., 2014].

In the near-field, the ball and stick neuron is relatively insensitive to the frequency of the ES up to about 100 Hz, decaying as 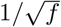 at high frequencies (Figure 4B-G, top row). Far away, the response is frequency-independent for low frequencies, eventually decaying as 1*/f* (Figure 4B-G, bottom row).

ES is relatively insensitive to changes in the neuron’s electrical properties (Figure 4B-D): in particular, the value of *R*_m_ is largely irrelevant. This may seem surprising: The somatic *V*_m_ response to current input is highly *R*_m_-dependent as can be understood from Ohm’s law, *V* = *RI. V*_e_, however, is fully determined by the membrane current; for a fixed current input, *R*_m_ therefore only plays a secondary role in the spatial distribution of the return currents.

For near-by ES, the length of the dendrite *l* is unimportant, because *V*_e_ is dominated by the somatic current alone. The effect of the dendritic length is more pronounced for distant ES (Figure 4E), since the length of the stick will directly affect the current-dipole moment through affecting the distance between the sinks and sources [Pettersen and Einevoll, 2008]. Interestingly, these results indicate that the spatial extent of neurons is primarily important for distant, low-frequency ES. This is also easily understood by considering the current-dipole moment for somatic current input: For high frequencies, the frequency-dependent length constant [Koch, 1999], which determines the spatial distribution of return currents, becomes much smaller than the total cell length, rendering the total length irrelevant at these frequencies.

The effect of ES is most sensitive to the diameter of the dendrite *d* (Figure 4F-G). For an increased *d*, the effect of ES increases (Figure 4F), as expected from the literature where it has been reported that large-diameter axons are most sensitive to ES [McNeal, 1976; Basser and Roth, 2000; Aberra et al., 2018; Ye et al., 2022]. The opposite is true for soma size *d*_s_, where the effect of ES decreases with an increased diameter (Figure 4G).

The opposite effect of the somatic and dendritic diameter is hard to make sense of through the traditional way of thinking about ES. However, through the RT-based approach it makes sense: We know that for somatic current input, a bigger soma will lead to more local and less distributed return currents, thereby resulting in a weaker current dipole moment and a lower amplitude *V*_e_. Through the RT we therefore know that a larger soma will decrease the sensitivity of the somatic *V*_m_ to ES. A thicker apical dendrite will have the opposite effect of less local and more distributed return currents, thereby resulting in a stronger current dipole moment.

This illustrates that in the traditional approach to simulating ES it is hard to build an intuitive understanding of how any neural property will affect how a neuron is affected by ES, but that the RT-based approach offers a new perspective on ES which can be helpful in this regard.

### 2.4. More complex extracellular mediums

Thus, the RT-based approach is helpful in analyzing the effect of ES on different neural elements embedded in an infinite, homogeneous, and isotropic medium. Of course, real brains are none of those [Wolters et al., 2006; Logothetis et al., 2007; Vorwerk et al., 2014]. However, as long as nervous tissue behaves approximately linearly, that is, varying the applied current varies the induced electrical field proportionally, the RT remains relevant, including when the extracellular conductivity is inhomogeneous (position-dependent), anisotropic (direction-dependent), or frequency-dependent.

For extracellular mediums with anisotropies or simple planar inhomogeneities in the extracellular conductivity, only minor changes are needed to the formalism for calculating *V*_e_ [Ness et al., 2015; Halnes et al., 2024]. Taking such effects into account is easy with appropriate simulators like LFPy [Hagen et al., 2018].

Cortical tissue is moderately anisotropic and inhomogeneous [Goto et al., 2010]; however, it has been shown that the effect on intracortical potentials is negligible [Gold et al., 2006; Logothetis et al., 2007; Ness et al., 2015], implying that we are justified in disregarding such effects for intracortical ES as well.

For measurements of potentials at the cortical surface (ECoG), the membranes and external materials covering the cortex can have a substantial effect on the recorded potentials [Pettersen et al., 2006; Ness et al., 2015; Rogers et al., 2020]: Electrically insulating materials (like non-conducting mineral oils or air) amplify signal amplitudes, while highly-conducting materials (like saline or metal) reduce signal amplitudes [Pettersen et al., 2006; Rogers et al., 2020; Halnes et al., 2024]. ES at the cortical surface will therefore have a larger effect on neural activity if a non-conducting material is used as a cover material.

It can be important to take the presence of the recording/stimulating equipment on extracellular potentials into account, in particular if it accounts for a large non-conducting volume that affects *V*_e_ [Moffitt and McIntyre, 2005; Lempka and McIntyre, 2013; Ness et al., 2015; Maling et al., 2018; Buccino et al., 2019; Bower and McIntyre, 2020; Molina-Martínez et al., 2022]. This necessitates numerical methods like the finite element method; yet, this does not invalidate the RT. Thus, Buccino et al. [2019] demonstrated that while microwires hardly influenced *V*_e_, the presence of the much larger Neuropixels and Neuronexus MEA probes boosted *V*_e_ from neurons in front of them by almost a factor of two, and decreased the signal amplitude from neurons behind them by about a factor of two (a shadowing effect). If the recording electrodes are instead used to inject currents, we can expect similar conclusions for the resulting membrane potential of nearby neurons.

The examples in the previous paragraphs highlights how the RT-based approach supplies a fresh perspective on ES, allowing us to take advantage of earlier research and insights.

### 2.5. Transcranial Electric Stimulation (tES)

Electroencephalography (EEG) signals, recorded on the scalp, are strongly influenced by the variable conductivities of cortical grey matter, cerebrospinal fluid, skull, scalp, and musculature [Wolters et al., 2006; Vorwerk et al., 2014; Nunez et al., 2019; Ness et al., 2022; Tharayil et al., 2024]. EEG signal prediction therefore warrants complex volume-conductor models, called *head models*. Importantly, all such models are linear [Nunez and Srinivasan, 2006; Ilmoniemi and Sarvas, 2019]. This enables us to apply the RT to infer changes in *V*_m_ responding to tES, that is, current stimulation from the scalp: we use a head model to calculate the EEG signal at a given location on the head (*V*_e_) due to a somatic current input to a neuron (operating in an approximately linear regime) at a given location in the brain. Through the RT, this corresponds to *V*_m_ of that neuron from tES at the EEG electrode location.

For calculating EEG signals from neural activity, it is common to represent neural activity as equivalent current dipoles. Calculating these from simulated neural activity is straightforward for multi-compartmental models [Hagen et al., 2018]. Furthermore, the link between current dipoles and the resulting EEG signals is well-developed [Nunez and Srinivasan, 2006; Ilmoniemi and Sarvas, 2019; Halnes et al., 2024], and current dipoles can be used with simple or detailed head models [Næss et al., 2021]. The EEG signal generated by a single current dipole **P**_*n*_(*t*) can, in general, be expressed as *V*_e_(**r**, *t*) = **M**(**r, r**_*n*_)**P**_*n*_(*t*), where **M** is the so-called lead-field matrix.

To illustrate the application of the RT to tES, we simulate a 10 Hz, 1 nA somatic current input to a passive L5 pyramidal cell model and calculate a resulting current-dipole moment amplitude of 186 nAµm (Figure 5A-C). Note that we here only consider the *z*-component, set to be along the apical dendrite, perpendicular to the cortical surface. The rationale is that because of rotational symmetry for pyramidal cells around the *z*-axis, the other components of the current-dipole moment will cancel for a population of pyramidal cells.

**Figure 5:**
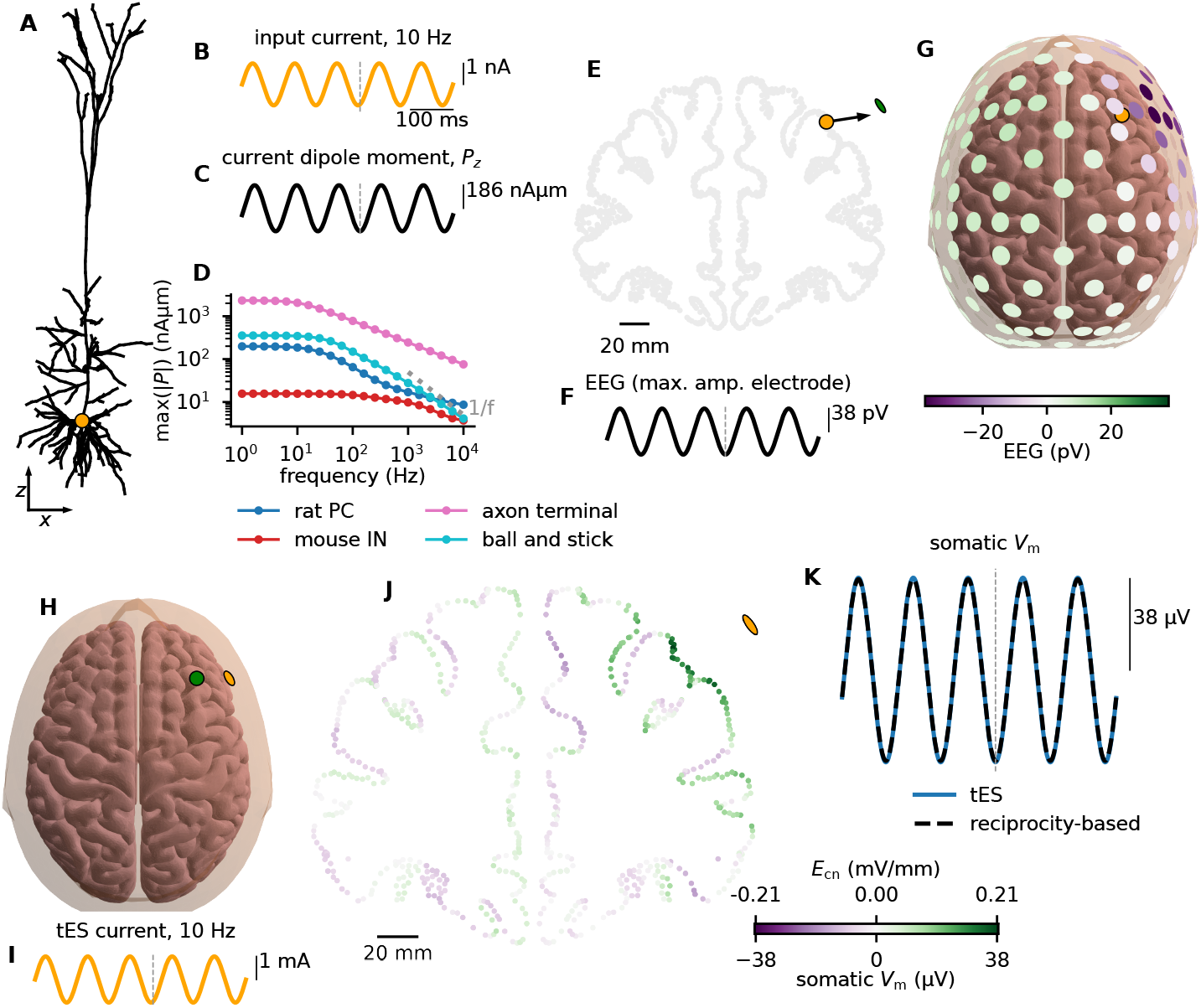
The effect of transcranial electric stimulation (tES) on individual pyramidal cells. **A:** A passive pyramidal cell model receiving a somatic current input (orange dot). **B:** The input current with a frequency of 10 Hz and a 1 nA amplitude. **C:** The *z*-component of the resulting current-dipole moment, perpendicular to the cortical surface. **D:** The amplitude of this *z*-component as a function of the input frequency for different cell models. **E:** The current-dipole (panel C) is placed in the middle frontal gyrus of the New York head model. The orange dot marks the location, the black arrow marks the orientation, and the green dot marks the nearest EEG electrode. **F:** The induced voltage signal of 38 pV at the nearest EEG electrode (green dot in panel E), from the single pyramidal cell at the location marked in panel E. **G**: The EEG signal at the time of the maximum signal (time indicated by gray dashed lines in panels B, C, and F). The location of the dipole is marked by the orange dot (partially covered by an EEG electrode). **H:** Through the RT we can invert the situation and calculate the somatic *V*_m_ for the pyramidal neuron (located at the green dot) responding to tES (located at the orange dot). **I:** A 10 Hz, 1 mA current input through an EEG electrode (marked in panel E and H), that is, 10^6^ times larger amplitude than the original intracellular current input. **J:** A cross section of the head model’s lead field, displaying the amplitude of the electric field along the cortical normal direction (*E*_cn_). For a tES current amplitude of 1 mA through the considered electrode (orange dot), the maximum value of *E*_cn_ is 0.21 mV/mm. This is the unmodified lead field as given by the New York head model [Huang et al., 2016]. The somatic *V*_m_ response for the cell model in panel A at different locations in the cortex, in response to the tES in panels H and I, can be found by a scaling of the lead field, in this case by a factor of 186 µm. **K:** The somatic *V*_m_ response to tES for the cell model in panel A at the location marked in panel E is 10^6^ · 38 pA = 38 µV. As a control, we also simulate the effect of a uniform electric field of amplitude 0.21 mV/mm along the *z*-axis of the cell model, and see that the somatic *V*_m_ response is indistinguishable.

We also quantify the relationship between the frequency of the input current and the resulting current-dipole moment amplitude for four different cell models (Figure 5D). For a ball and stick model, the current-dipole moment is frequency-independent for low frequencies, falling off as 1/*f* at higher frequencies (Figure 4), while the decay is somewhat less steep for the other three models. For the pyramidal cell, the amplitude decays from 198 nAµm at 1 Hz to 8.6 nAµm at 10 kHz, that is, by a factor of 23 for a frequency increase over 4 orders of magnitude. The axon terminal is about an order of magnitude more sensitive to tES than the pyramidal cell, consistent with previous simulation results [Aberra et al., 2023].

To calculate the signal on the scalp evoked by this particular neuron, we use the New York head model, as it is a detailed model with a freely available lead field for each EEG electrode. The amplitude of the evoked potential at a simulated EEG electrode above the middle frontal gyrus (Figure 5E) is 38 pV (Figure 5F), decaying with distance from the dipole, and changing sign, as expected from a dipole (Figure 5G). Of course, such an tiny evoked signal would not be detectable, given the ambient biological and instrumental noise levels, but constitutes an idealized estimate.

Via the RT, this implies that a 10 Hz, 1 nA current injection through the same EEG electrode will induce 38 pV change in *V*_m_. Common current amplitudes for tES are on the order of 1 mA [Dmochowski et al., 2017; Sliva et al., 2018; Liu et al., 2018; Krause et al., 2019], a million times bigger. That is, a 10 Hz, 1 mA tES stimulus will induce a *V*_m_ amplitude of 38 pV × 10^6^ = 38 µV (Figure 5H-K). A million-fold scaling factor might at first glance seem questionable, but note that the only relevant constrain is that the resulting *V*_m_ should be subthreshold, and 38 µV remains well within the linear regime (Figure 2E). Even a ten times larger current of 10 mA, above the safety level for human usage [Antal et al., 2017], would induce an evoked *V*_m_ of only a third of a millivolt.

The example in the previous paragraph treated a single cortical location and many EEG electrodes, but it is equally possible to focus on a single electrode and evaluate the *V*_m_ response for the given cell model at any cortical site. For the somatic 10 Hz, 1 nA input to the pyramidal cell model, the amplitude of the resulting current-dipole moment was 186 nAµm. By multiplying this number with the lead field **M**(**r, r**_*n*_), and further scaling it by 10^6^, we obtain the subthreshold somatic *V*_m_ response at any cortical location (Figure 5J).

Cortical regions where the normal points towards the electrode typically experience the opposite effect relative to regions facing away from the electrode (Figure 5J). As expected, tES is strongest close to the electrode, but overall its effect is quite evenly distributed across the brain. Although the amplitude will be frequency dependent (Figure 5D), the spatial distribution will not be, since this term comes solely from the head model, which is frequency-independent [Ranta et al., 2017; Huang et al., 2017; Vöröslakos et al., 2018].

Note that the lead field **M**(**r, r**_*n*_) can be interpreted as the electric field resulting from tES [Rush and Driscoll, 1969; Huang et al., 2016; Dmochowski et al., 2017]; the amplitude of this field along the cortical normal direction (*E*_cn_), is related to the amplitude of the somatic *V*_m_ response through a (frequency-dependent) conversion factor (Figure 5J, two different color scales).

As a final control, we directly simulate the somatic *V*_m_ response of the considered neuron model (Figure 5A) under the influence of the uniform electric field given by the head model (Figure 5J) at the previously used cortical location (Figure 5E). This approach assumes that the electric field is relatively uniform over the spatial extent of the neurons, and is sometimes called the quasi-uniform assumption [Bikson et al., 2013; Aberra et al., 2023]. As for the infinite homogeneous medium (Figure 2), the direct simulation of the effect of tES (Figure 5K, blue line) and the reciprocity-based approach (Figure 5K, dashed black line) yield indistinguishable results.

## 3. Discussion

We here use the reciprocity theorem (RT) to shed light on the neural response to extracellular current stimulation (ES) (Figure 1). No error is introduced using RT to estimate the effect of ES on passive cell models; errors are minimal (a few percent) for active cell models in the relevant regime, which is the subthreshold one (Figure 2). We use the RT-based approach to study how different neural elements are affected by ES (Figure 3). Furthermore we derive an analytic expression for the somatic *V*_m_ response of a passive ball and stick model under the influence of ES, extracting simple power laws for its dependency on frequency and distance, and comparing the sensitivity of ES for different neural parameters (Figure 4). Finally, we demonstrate that the RT is applicable to study transcranial electric stimulation (tES): Using a detailed head model we can directly predict the subthreshold *V*_m_ response to tES of an arbitrary cell model at any location in the brain (Figure 5).

### 3.1. What is new?

The RT has a long history in neuroscience, yet its ability to directly study ES has been overlooked: It has so far only been used to swap the locations of extracellular current sources (or current dipoles), and extracellularly measured potentials. We here demonstrate that applying the RT to study ES has many advantages: We show that the RT explains the surprising cell-type and frequency-independent ES results presented by Lee et al. (2024), and nuances these results by demonstrating that they are only expected to hold for nearby neurons below about 100 Hz. Furthermore, the RT-based approach supplies us with analytic formulas and power laws for both the frequency-response and distance-dependence of ES, which has not, to the best of our knowledge, been reported previously.

In general, moving from intracellular stimulation of a neuron to predicting the resultant *V*_e_ at *n* locations to injecting an extracellular current, as in transcranial ES, and estimating the effect on *V*_m_ at *n* location, has the same computational complexity in terms of number and order of equations to be solved. The reason for why the RT provides a fast and intuitive way of understanding ES, is that electrophysiology has a biased view of neurons, emphasizing the location and size of the cell body, at the expense of other neuronal regions: the soma is far more accessible to intracellular stimulation and recording than other neuronal compartments and is in close proximity of the axon initial segment where *V*_m_ is transduced into one or more spikes. This soma-centric view has led to the development of a mature understanding regarding the origin of extracellular potentials following somatic current injections. The RT also supplies a new perspective on ES which makes it easier to understand how different neural features, like morphology and diameters, influences a neuron’s susceptibility to ES. We therefore conclude that the RT-based approach is capable of providing new general insights into how ES affects neurons.

### 3.2. When to apply the RT-based approach for studying ES

The RT allows us to rely on the extensive knowledge available on how neural activity generates extracellular potentials [Halnes et al., 2024], to gain a better intuition of how ES works, instead of the traditional approach which involves explicitly simulating each specific combination of stimuli and cell model, including the locations and time course of the stimuli. In the RT-based approach, a single simulation of a somatic white noise current input with simultaneous calculation of *V*_e_ at all relevant extracellular locations, is sufficient to obtain the subthreshold response of that particular cell model to an ES at any one of the considered locations. Furthermore, since the system is linear and we have the full frequency response, we can, through Fourier analysis, find the subthreshold somatic *V*_m_ response to an arbitrary time course of the ES, including for mono- or bi-phasic pulses of arbitrary durations.

We here considered the effect of ES on one neuron; intriguingly, our results show that the linear *V*_m_ response of a set of *N*_*n*_ neurons at different locations, resulting from a set of *N*_*i*_ stimulation electrodes can be formulated as a matrix equation in frequency space **V**_m_ = **MI**_stim_, where **M** is the lead field which through the RT can be found by computing extracellular potentials due to current stimulation of the soma. We can tailor **I**_stim_ towards a desired **V**_m_ response in the *N*_*n*_ neurons.

### 3.3. Transcranial electric stimulation

Our results indicate that tES in the range of 1-4 mA on the scalp induces membrane potential changes of 38-152 µV at the soma of a single pyramidal cell inside the human brain. This is consistent with previous findings: 1-4 mA tES gives a 0.21 - 0.84 mV/mm field in the human brain for the head model we use, and similar values are reported by others [Vöröslakos et al., 2018]. In rat hippocampus, somatic depolarizations on the order of 0.2 mV per V/m of applied field have been observed [Bikson et al., 2004; Reato et al., 2010; Liu et al., 2018]. For the field values listed above (0.21 - 0.84 mV/mm), this corresponds to 42-168 µV somatic depolarization, in line with our results.

Grossman et al. [2017] directly induced spikes in mice with current amplitudes of ca. 0.5 mA. Yet the same current amplitudes would give two orders of magnitude lower electric field amplitudes in the human brain [Alekseichuk et al., 2019]. This is consistent with predictions via the RT: Halnes et al. [2024] showed that EEG signals in a mouse head model were two orders of magnitude larger than EEG signals in a human head model from the same neural source. The primary reason is the 10-fold larger distance between the scalp and neural sources, translating into a 100-fold smaller *V*_e_ in people and mice (based on the dipole decay of 1*/r*^2^; Figure 4A). Increasing currents by a 100-fold to compensate this signal loss is problematic; indeed, such large currents were historically employed to induce electric anesthesia [Liu et al., 2018; Vieira et al., 2024]. It therefore seems likely that observed effects of tES in humans must come about indirectly by either enhancing the synchrony of firing of specific neuronal populations in specific frequency ranges or from mechanisms like stochastic resonance [Liu et al., 2018], as suggested by Caldas-Martinez et al. [2024].

It was previously suggested, as a rule of thumb, that an electric field of 1 mV/mm is sufficient to entrain neural activity [Ozen et al., 2010; Anastassiou et al., 2011; Vöröslakos et al., 2018; Cakan and Obermayer, 2020]; however, neural effects of field strengths as low as 0.2 mV/mm are well documented [Reato et al., 2010; Liu et al., 2018; Johnson et al., 2020; Tran et al., 2022; Alekseichuk et al., 2022; Zhao et al., 2024]. In the detailed New York head model applied here, a 1 mA tES yields electric fields of 0.2 mV/mm (Figure 5J). tES amplitudes of 1-5 mA should therefore be in the lower range of what has been reported to affect human neural activity. We estimate that this correspond to somatic membrane potential responses on the order of 40-200 µV at 10 Hz, consistent with previous results from both simulations and experiments [Bikson et al., 2004; Reato et al., 2010; Aberra et al., 2023].

That the effect of tES on membrane potentials is much smaller than the spiking threshold has implications for how tES is mediated: Axon terminals were here found to be about an order of magnitude more sensitive to ES than pyramidal cells (Figure 3, Figure 5, Aberra et al. [2023]); however, the membrane potential of axon terminals are expected to lie stably around rest (when not firing an action potential), and are therefore less likely to be influenced by minor induced changes in their membrane potential than somas, where the membrane potential occasionally fluctuates close to the firing threshold. Therefore, while axons are likely to be the main target of intracranial microstimulation [Nowak and Bullier, 1998; Histed et al., 2009; Aberra et al., 2018], it seems more likely that the spike initiation zone is the main target of tES.

### 3.4. Limitations

The principal limitation in using the RT-based approach to simulate the effect of ES is the underlying assumption of linearity, which must hold for both the neuron and the extracellular environment (brain and head). Most studies report that the electrical properties of the brain and head are approximately linear and ohmic for frequencies below several kHz [Pfurtscheller and Cooper, 1975; Logothetis et al., 2007; Miceli et al., 2017; Ranta et al., 2017; Huang et al., 2017; Vöröslakos et al., 2018; Lee et al., 2024]. Some studies report weakly frequency-dependent tissue [Wagner et al., 2014; Vieira et al., 2024]; for the high frequencies of temporal interference stimulation, it is plausible that such effects can become important and might challenge the quasi-static assumption [Bossetti et al., 2008]. Note, however, that a frequency-dependent head model would still typically be linear [Nunez and Srinivasan, 2006; Halnes et al., 2024], and as such, the assumption of linearity with regards to the extracellular environment seems well-justified.

Neurons, on the other hand, have non-linear membrane properties, in particular when operating close to their spiking threshold. Mirzakhalili et al. [2020] demonstrated how an ion-channel-mediated rectification mechanism could explain how neurons respond to the envelope frequency during TI stimulation. It is plausible that such non-linear effects are needed to explain how spikes are directly evoked by extremely high-amplitude temporal interference stimulation, such as described by Grossman et al. [2017] (see also discussion in Vieira et al. [2024]). However, it is not obvious that this particular kind of non-linear ion-channel effects are critical for understanding the network mechanisms underlying the effect of TI in humans, such as stochastic resonance [Liu et al., 2018; Caldas-Martinez et al., 2024; Vieira et al., 2024]. Indeed, how weak *V*_m_ perturbations affect neural networks might be easiest to understand through simplified network simulations (see Outlook).

Although our results here indicate that the tested neuron models behaved close to linearly in the subthreshold regime (Figure 2), some ion channels are known to be active also in the subthreshold regime. In particular, *I*_h_ affects both extracellular potentials [Ness et al., 2016, 2018] and how neurons are affected by weak electric fields [Aspart et al., 2018]. However, we can expect *I*_h_ to primarily be important for *V*_m_ in the apical dendrite. Furthermore, calcium spikes are probably more likely to be evoked when apical dendrites are depolarized [Huang et al., 2024]. Therefore, in certain cases, the assumption of linearity might not be appropriate.

### 3.5. Outlook

Ours and previous results indicate that the direct effect of tES on the membrane potential of individual neurons is weak, in the range of ten to a few hundreds of µV, far away from the threshold for spiking. This implies that any behavioral and cognitive effects of TI must be mediated by synergistic network effects, such as by stochastic resonance and/or by advancing or delaying spike times [Reato et al., 2010; Liu et al., 2018; Krause et al., 2023; Caldas-Martinez et al., 2024].

A common approach to studying network effects is via point-neuron networks or mean-field models, and this is also true for studying network effects from tES [Aspart et al., 2016; Ladenbauer and Obermayer, 2019; Duchet et al., 2020; Cakan and Obermayer, 2020; West et al., 2022; Zhao et al., 2024], where better models for brain dynamics could aid understanding tES [Krause et al., 2023].

The RT-based approach is an alternative way of estimating the direct effect of tES, which can be implemented in such models. Note that we do not claim that the effect of tES on neural dynamics is linear, but that the direct, isolated effect of tES on membrane potentials usually is: Also in the traditional way of simulating, the ES enters as an additive term in the cable equation, (1*/r*_i_)∂*E*_||_(*x, t*)*/*∂*x*, which only depends on *E*_||_ and the cable properties, and not on the neural membrane potential. The ES-induced effect itself on the membrane currents is therefore linear and additive, even when the resulting membrane potential dynamics is not. Therefore, if we use the RT-based approach to estimate the somatic membrane potential amplitude response to tES, this could be added to the membrane potential of point neurons in a network model. The resulting neural network dynamics can be non-linear.

The assumption of linearity is not so restrictive at it seems, as it incorporates all active currents, as well as the detailed cellular morphology. This approach will therefore be more accurate and efficient than estimations using other approximations.

The RT-based approach can directly predict the somatic *V*_m_ effect from ES throughout a volume, for different cell types, and under different stimulation parameters (see also Gaugain et al. [2024] for a similar approach), which can be incorporated in point-neuron network simulations using NEST [Gewaltig and Diesmann, 2007], or even whole-brain mean-field models using *The Virtual Brain* [Leon et al., 2013].

## 4. Methods

### 4.1. Neural simulations

All simulation code was written in Python, and neural simulations were done through LFPy 2.3 [Hagen et al., 2018] running on NEURON 8.1 [Carnevale and Hines, 2006].

### 4.2. Cell models

The pyramidal cell model that was used was the rat cortical layer 5 cell model presented by Hay et al. [2011]. The interneuron model was a perisomatic layer 5 aspiny PV cell from the mouse primary visual area from the Allen Brain Atlas (celltypes.brain-map.org/; electrophysiology ID: 469610831; model ID: 491623973). The parameters for the myelinated axon model were from Hallermann et al. [2012], in the implementation used by Thunemann et al. [2022]. The total length was 10 mm, and a node of Ranvier was inserted for every 50 µm. We also confirmed that results were similar for the myelinated axon model created by McIntyre et al. [2002]. The axon terminal was just an axon endpoint with the same diameter as the rest of the axon.

Passive versions of the cell models were modified versions of the original cell models, where all active conductances were removed.

### 4.3. Input to cell models

Intracellular input currents were modeled as synapse currents (“POINT_ PROCESS” in NEURON), that is, they are treated as membrane currents. This is because the membrane potential is measured across the membrane (“port 1” in Figure 6A), and when applying the RT, the current should also be across the membrane (“port 1” in Figure 6B).

**Figure 6:**
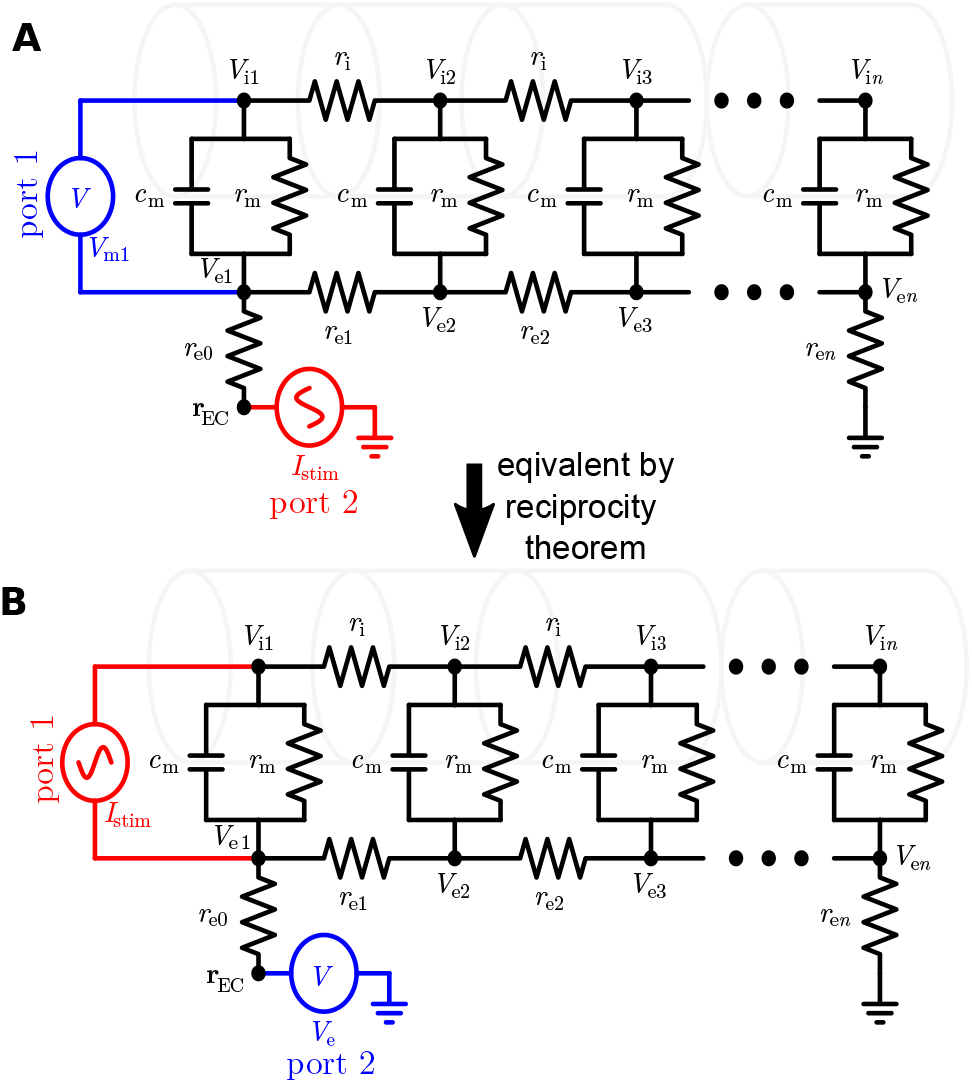
Illustration of the reciprocity theorem. **A:** The equivalent circuit diagram for a passive neuron model stimulated by an extracellular current source *I*_EC_, while the membrane potential of compartment 1 is measured. **B:** The equivalent circuit diagram for a passive neuron model stimulated by an intracellular current source *I*_EC_, while the extracellular potential is measured. According to the reciprocity theorem, the measured potential in the two

For intracellular current input to active cell models (Figure 2), we relied on a hand-picked, fixed cell-specific current amplitude in the subthreshold regime. This was done because the different cell models have vastly different input impedances, and react differently to the same current input. The amplitude was 0.1 nA, 0.01 nA, and 0.001 nA for the pyramidal cell model, interneuron model, and axon model, respectively. The resulting extracellular potential was afterwards scaled to reflect the amplitudes of the extracellular current source.

White-noise current input was constructed as a sum of sinusoids, where each frequency component had the same amplitude (1 nA) and a random phase [Lindén et al., 2010; Ness et al., 2016; Miceli et al., 2017]. The frequency components used when constructing the white noise were the same as those considered when extracting amplitude- and phase spectrums through Fourier analysis.

### 4.4. Simulating extracellular stimulation

The effect of an external extracellular potential on a neuron can be simulated with LFPy [Hagen et al., 2018], running on top of the NEURON simulation environment [Carnevale and Hines, 2006].

#### 4.4.1. Infinite homogeneous mediums

The extracellular potential at the midpoint location of each neural compartment is calculated for each time step, and inserted into NEURON as a boundary condition through the extracellular mechanism. For infinite homogeneous mediums, the extracellular potential from an ES point source *I*_stim_(*t*) at location **r**′ can be calculated as [Halnes et al., 2024],

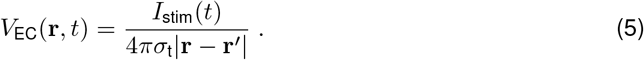

#### 4.4.2. Detailed head model

Since the tES electrodes are far away relative to the size of neurons, we use the quasi-uniform assumption [Bikson et al., 2013; Aberra et al., 2023] that the electric field is relatively constant over the extent of the neuron, and calculate the resulting extracellular potential along the *z*-axis, as *V*_EC_ = *E*_cn_*z*, where *E*_cn_ is the electric field along the cortical normal direction (the *z*-axis of the local coordinate system), which we obtain directly from the lead field matrix of the New York head model [Rush and Driscoll, 1969; Huang et al., 2016] (https://www.parralab.org/nyhead/).

### 4.5. Signal analysis

For the Fourier analysis we used the `scipy.fftpack` Python package to extract the amplitude *V*_e_(*f*) and phase *θ*(*f*) spectrums from calculated signals. When plotting the spatial profiles of extracellular potentials at specific frequencies (Figure 3), the phases of *V*_e_ was incorporated through plotting Re{*V*_e_(*f*)*e*^*iθ*(*f*)^}.

### 4.6. Analytic expressions

For the somatic membrane current, the transfer function is given by [Pettersen et al., 2014]

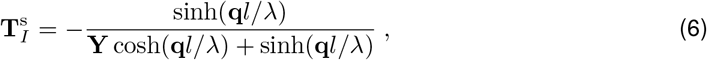

and for the current-dipole moment the transfer function is given by,

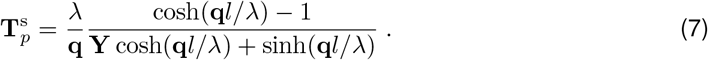

Here 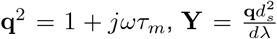, and *l* and *d* is the length and diameter of the stick, respectively, while the diameter of the soma is *d*_*s*_.

### 4.7. Application of the reciprocity theorem

One can only expect the RT to be applicable for approximately linear systems, which includes passive cell models, that is, cell models without any active conductances, and active cell models which are operating in the subthreshold regime, where neurons often behave approximately linearly [Koch, 1984; Bikson et al., 2004; Remme and Rinzel, 2011; Ness et al., 2016, 2018; Gaugain et al., 2024]. However, the amplitude of extracellular current sources used to stimulate neurons (1000 nA in Figure 2B) is a typical amplitude that would drive neurons to fire action potentials if given intracellularly, and the neurons would therefore be far outside the approximately linear regime. Directly using the same current amplitudes for extracellular and intracellular current sources is therefore applicable for passive cell models, but not for active cell models. For intracellular current input to active cell models (Figure 2) we therefore relied on the assumed approximate linearity in the subthreshold regime and used a fixed, cell-specific, subthreshold current amplitude (0.1 nA for the pyramidal cell model in Figure 2A). The resulting extracellular potential were afterwards scaled by the ratio of the extracellular to intracellular current amplitude (that is, by a factor of 1000 nA / 0.1 nA = 10,000 in Figure 2B).

In electromagnetism, the reciprocity theorem states that for time-invariant linear media the relationship between an oscillating current and the resulting electric field is unchanged if one interchanges the points where the current is placed and where the field is measured [Hayt et al., 2019]. When simulating neural activity, it is standard practice to represent neurons as equivalent electric circuits [Hodgkin and Huxley, 1952; Rall and Shepherd, 1968; Koch, 1999]. For the special case of electric circuits, the reciprocity theorem states that in any passive linear network, if a current source *I*_0_ between nodes *x* and *x*′ produces the voltage response *V*_*y*_ between nodes *y* and *y*′, then the removal of the current source from nodes *x* and *x*′ and its insertion between nodes *y* and *y*′ will produce the voltage response *V*_*y*_ between nodes *x* and *x*′ [Hayt et al., 2019].

In the context of electric stimulation of neurons, this implies that if there is a current source *I*_stim_ at extracellular location **r**_EC_, flowing to ground (Figure 6A “port 2”), and this produces the voltage response *V*_rec_ over a given part of a neural membrane (Figure 6A “port 1”), then moving the current source to instead be across the neural membrane (Figure 6B “port 1”) will produce the same voltage response *V*_rec_ between location **r**_EC_ and ground [Hayt et al., 2019] (Figure 6B “port 2”).

The reciprocity theorem is valid for any electrical network that consists entirely of ideal capacitances, inductances, and resistances, that is, passive electrical networks. For simplicity, one can start by analyzing the electrical network in the absence of the resting membrane potential, because the principle of superposition ensures that the resting membrane potential can be added afterwards as additional voltage sources (“batteries”) in the equivalent circuit. When the resting membrane potential is not taken into account, the RT-based approach only predicts deviations of *V*_m_ from the resting membrane potential.

Because of the conventions regarding the direction of membrane currents, a sign change is needed for the intracellular current input (relative to the extracellular current source) when it is inserted in NEURON as a synapse current.

### 4.8. Code availability

All simulation code is freely available through https://github.com/torbjone/ES_from_reciprocity.

## Acknowledgements

T.V.N. and G.T.E. received funding from the European Union Horizon 2020 Research and Innovation Programme under Grant Agreement No. 101147319 [EBRAINS 2.0]. C.K. thanks the Allen Institute founder, Paul G. Allen, for his vision, encouragement, and support.

